# Unimodal effects of pigment richness on niche and fitness differences explain species richness and ecosystem function in light-limited phytoplankton communities

**DOI:** 10.1101/2020.07.28.225003

**Authors:** Jurg Werner Spaak, Frederik De Laender

**Affiliations:** Research Unit in Environmental and Evolutionary Biology, Institute of Life, Earth and the Environment, and Namur Institute of Complex Systems, University of Namur, Belgium.

**Keywords:** coexistence, ecosystem function, niche differences, phyto-plankton, pigment, species richness, trait diversity

## Abstract

Trait diversity is traditionally seen as promoting species richness and ecosystem function. Species with dissimilar traits would partition available resources, increasing niche differences, facilitating coexistence and increasing ecosystem function. Here we first show, using theory and simulations for light-limited phytoplankton, that combing photosynthetic pigments is indeed a necessary condition for coexistence and stimulates ecosystem function. However, pigment richness does mostly not permit the coexistence of more than two species, and increases productivity at most 60% compared to single-pigment communities. Surprisingly, combining all nine pigments known to date leads to a 2.5% probability that four species would coexist, illustrating that the coexistence of a high number of species along a continuous niche axis is constrained by limiting similarity. We explain these constraints by unimodal effects of pigment richness on niche and fitness differences, which jointly limit the positive effect of pigment on species richness. Empirical data and additional simulations suggest that pigment richness effects can be stronger during transient dynamics but inevitably weaken with time, i.e. pigment richness effects on species richness and function are likely short-lived. Our results highlight the need to apply coexistence theory to understand the long-term effects of trait diversity on biodiversity and ecosystem function.

**Statement of authorship:** J.W.S. and F.dL. developed the ideas and wrote the manuscript. J.W.S developed the mathematics and the python code to conduct the study. J.W.S conducted the literature review.

## Introduction

Species richness is a main predictor of ecosystem function and stability (Hector & Loreau 2001; Hooper *et al.* 2005 but see Spaak *et al.* 2017; Srivastava & Vellend 2005). Communities with more species typically produce more biomass and more stably do so than communities with few species (Striebel *et al.*, 2009; Tilman, 1996; Hector & Loreau, 2001; Balvanera *et al.*, 2006). Thus, identifying the factors that sustain and limit the capacity of species to coexist is an essential task, and probably one of the most fundamental objectives in ecology (Chase & Leibold, 2003; Hubell, 2001; Chesson *et al.*, 2001; Tilman, 1982).

In the past three decades of community ecology, traits have been considered a key ingredient to explain species coexistence (Mcgill *et al.*, 2006; HilleRisLambers *et al.*, 2012). Traits determine both how species respond to environmental variation and to the presence of other species (Chase & Leibold, 2003) and it is generally accepted that trait diversity drives niche differences, which act to stabilize coexistence (McKane *et al.*, 2002; Mayfield & Levine, 2010). Indeed, when all species have identical traits, trait diversity is zero and stable coexistence is not possible (Chesson, 2000; Bell, 2000; Hubell, 2001). In addition, trait diversity allows coexisting species to exploit a more diverse set of resources and do so more completely, optimising function.

Recent progress in coexistence theory has shown that trait diversity can have both positive and negative effects on biodiversity, including species richness. That is because, apart from increasing stabilizing niche differences, trait diversity can also increase fitness differences (HilleRisLambers *et al.*, 2012; Gallego *et al.*, 2019; Narwani *et al.*, 2013), which can act to disrupt coexistence when they overrule the stabilizing effects. Whether trait diversity supports or limits the number of species that can stably coexist, and thus biodiversity, is therefore unsure (Violle *et al.*, 2011; Best *et al.*, 2013; Levine & HilleRisLambers, 2009). Do species coexist be-cause they are sufficiently different or because they are sufficiently similar?

Here, we theoretically investigate the effects of trait diversity (the number of different traits, i.e. trait richness) on the number of stably coexisting species, niche and fitness differences and ecosystem function (total biovolume). Our working hypothesis was that a greater trait richness would lead to higher species richness and function because it would increase niche differences more than fitness differences, corresponding to prevailing ideas on the effects of trait richness. We considered light limited phytoplankton communities as a globally relevant study system that drives most aquatic food-webs. (Field *et al.*, 2009; Irigoien *et al.*, 2004; Striebel *et al.*, 2012; Baggio *et al.*, 2018; Kulkarni & De Laender, 2017).

Competition for light in phytoplankton communities is a major process explaining community composition throughout the world’s aquatic habitats (Goldman *et al.*, 1979; Langdon, 1988; Stomp *et al.*, 2007). Phytoplankton communities in meso- and eutrophic lakes and oceans, where light is the main limiting factor, produce approximately 30%-40% of the world’s annual primary production (Field *et al.*, 2009; Berger *et al.*, 2006; Boyd, 2002). In addition, competition for light is particularly suitable to test our expectations because the light spectrum represents a continuous resource axis, which a high number of species could in principle be able to partition.

The traits involved in competition for light are pigments and photosynthetic efficiency, i.e. the amount of biovolume gained per absorbed photon (Huisman & Weissing, 1994; Stomp *et al.*, 2004). Differences of pigmentation phenotypes have been shown to facilitate coexistence among, for example, cyanobacteria species (Stomp *et al.*, 2004, 2007). Pigment diversity is also expected to allow a more complete utilisation of the light spectrum (Striebel *et al.*, 2009), thus promoting both species richness and optimizing function. Photosynthetic efficiency alone cannot facilitate coexistence, the reason for which we do not consider this trait further.

We start this paper by theoretically examining how pigment richness determines the number of stably coexisting phytoplankton species and ecosystem function (total biovolume). We do so by analysing a phytoplankton growth model that incorporates the partitioning of the light spectrum among species with different pigments (Stomp *et al.*, 2004). We show that, when all species share one pigment, communities evolve to mono-dominance. When a community contains multiple of the main nine photosynthetically active pigments found in nature, up to four species co-exist based on light spectrum partitioning, but with low probability. We explain this result through unimodal effects of pigment richness on niche and fitness differences. Higher pigment richness also increases total biovolume by approximately 60%.

Next, we compile data from the literature and find a much stronger positive effects of pigment trait richness on species richness and ecosystem function. We reconcile these data with additional model simulations, finding that the reported effects are likely short-lived (transient) phenomena. Our findings highlight that applying coexistence theory is needed to understand the mechanisms linking species richness to ecosystem functions on time scales relevant for natural systems.

## Methods

### Model description

We used a model proposed earlier by Stomp *et al.* (2004), which is equivalent to (Appendix S1):

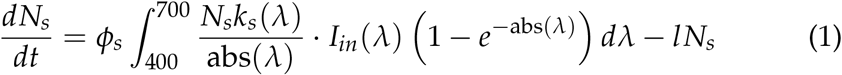

Where *N*_*s*_ is the density of species *s* (in *fl*/*ml*), *ϕ*_*s*_ is the photosynthetic efficiency, *k*_*s*_(*λ*) is the absorption spectrum of species *s*, 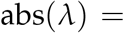 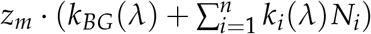 is the sum of all absorbers (background and phytoplankton species), where *k*_*BG*_ is the background absorption, *z*_*m*_ is the mixing depth and *n* is the species richness, *I*_*in*_(*λ*) is the incoming light intensity and *l* is the loss rate. Note that we considered the loss rate not as a species-specific parameter but rather the dilution rate of the system. The integral is taken over the whole range of photosynthetic active radiation (400nm - 700nm).

*I*_*in*_(*λ*)(1 − *e*^−abs(*λ*)^) can be thought of as the total amount of photons absorbed by the system and 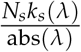 is the fraction of these photons that species *s* uses for its growth.

### Analyses and simulations

We used equation 1 to examine how trait (pigment) richness affects species richness and ecosystem function. We first theoretically analysed the maximum number of species that can stably coexist based on a predefined number of pigments.

Next, we ran simulations to determine the actual number of species and level of ecosystem function corresponding to a given pigment richness. To do so first, we collected 15 algal pigmentation types from the literature representing the major pigmentations of marine and freshwater phytoplankton (Six *et al.*, 2007; Van Den Hoek *et al.*, 1995) (Appendix S2). Then, we randomly assembled 400‘000 communities composed of a random number of species (1 to 15). In every community, every species was randomly assigned a pigmentation type. The absorption spectrum of a species is defined as the sum of it’s pigments:

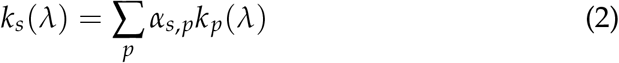

Where *k*_*s*_ is the absorption spectrum of the species, *k*_*p*_ is the absorption spectrum of the pigment and *α*_*s,p*_ is the concentration of pigment *p* in species *s*. The identity of the pigments present in a species is given by its pigmentation type. The absorption spectra of the pigments (*k*_*p*_(*λ*)) were taken from the literature (Bricaud *et al.*, 2004; Six *et al.*, 2007). Fig. 1 B,C shows an example for two pigmentation types. The total absorption of each species was kept constant 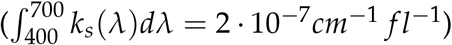, to avoid species with more pigments absorbing more light. Furthermore, we varied the photosynthetic efficiency 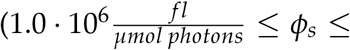 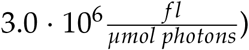 (Langdon, 1988), the dilution rate (0.003*h*^−1^ ≤ *l* ≤ 0.015*h*^−1^) (Stomp *et al.*, 2004; Striebel *et al.*, 2009) and the incoming light intensity (20*μmol photons m*^−2^*s*^−1^ *≤ I*_*tot*_ *≤* 200*μmol photons m*^−2^*s*^−1^) (Stomp *et al.*, 2004; Striebel *et al.*, 2009). We assumed that the background absorption is negligible (*k*_*BG*_ ≈ 0, Appendix 3 for simulation with non-zero background absorption).

**Figure 1:**
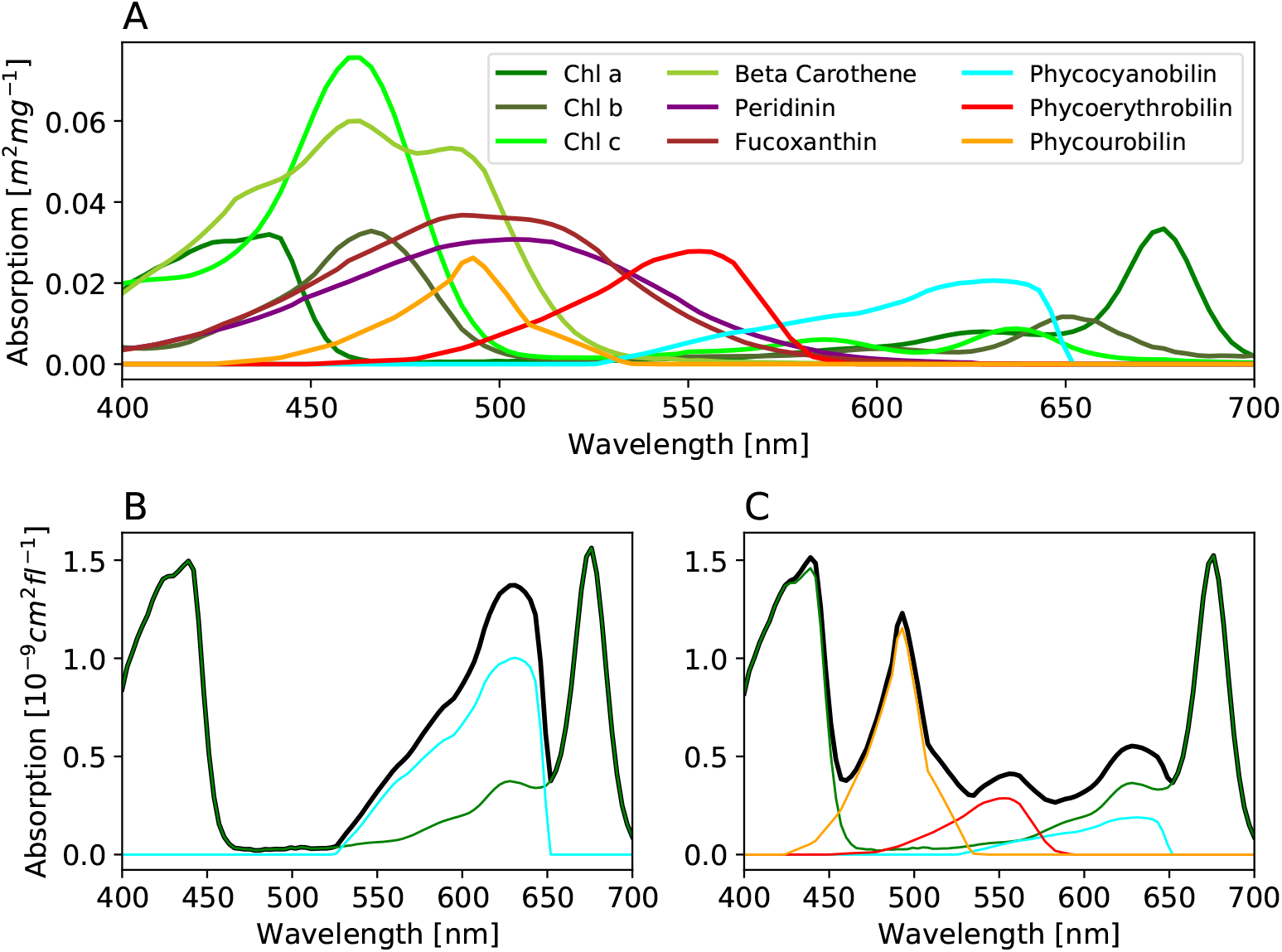
In vivo absorption spectra of all considered pigments (A) and two examples (B: Cyanobacteria type 1, C: Cyanobacteria type 3) showing how absorption spectra (black) can be decomposed into different pigments (Eq. 2). The two species in B and C can coexist under white light, as the Cyanobacteria type 1 is a superior absorber in the red light (620-700 nm), while the Cyanobacteria type 3 species is a better absorber in the blue-green light (450-550 nm).

For every community we measured species richness, pigment richness and ecosystem function over time. To measure species richness and pigment richness we assumed that species with relative densities below 0.01% are extinct. Changing this threshold to 1% or 0.0001% did not change our results. The ecosystem function we considered was total biovolume 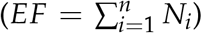. The ecosystem function was measured in communities with intensity of incoming light fixed at 40*μmol photons m*^−2^*s*^−1^. Again, changing this value to 20*μmol photons m*^−2^*s*^−1^ or 80*μmol photons m*^−2^*s*^−1^ did not qualitatively alter the results.

For each community at equilibrium we computed the niche and fitness differences (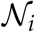 and 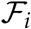) of the persisting species, using the definition Spaak & De Laender (2020). Technically, Spaak & De Laender (2020) define 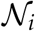 based a species *i*’s growth rate under various conditions, including its invasion growth rate (i.e. its growth rate when seeded in a community of other species) and its intrinsic niche growth rate (i.e. at low density in absence of competitors). As the difference between the invasion and intrinsic growth rate gets larger, 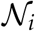 will approach zero, which indicates strong species interactions. In the current context, we expect this to correspond to the case where species have identical pigmentation, and thus compete maximally for light. Conversely, as the difference between the invasion and intrinsic growth rate gets smaller, 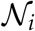 will approach one, which indicates weak species interactions. We expect this to correspond to the case where species differ markedly in pigmentation. Spaak & De Laender (2020) define 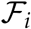 as the growth rate of species *i* in absence of niche differences, scaled by the intrinsic growth rate of species *i*. In that case, the species with the highest scaled growth rate has the largest fitness difference. Because these definitions do not lead to closed forms of 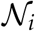 and 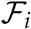, we computed 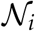 and 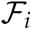 numerically.

### Literature data

We searched the literature for experimental data describing the effects of pigment richness on species richness and ecosystem function. We used the search term (phytoplankton OR algae) AND (“pigment richness” OR “pigment diversity”) AND (“ecosystem function” OR biodiversity) using the search engines scopus and google scholar. Additionally, we screened all papers cited by or citing as such identified papers. We identified four datasets (two representing field samples from freshwater lakes and two representing experiments with freshwater and marine phytoplankton respectively). The two field data sets represent different sampling locations in one lake (Fietz & Nicklisch, 2004) and from multiple lakes (Striebel, personal communications), respectively. The experimental dataset of Estrada *et al.* (2004) represents phytoplankton diversity in solar salterns with salinity ranging from 4% to 22.4%. We excluded salinities above 22.4%, as they do not resemble marine ecosystems (Estrada *et al.*, 2004). Striebel *et al.* (2009) assembled 1 to 10 species of freshwater phytoplankton and measured pigment richness and ecosystem function after 14 days. Pigment richness was measured using high-performance liquid chromatography in all datasets. The datasets contain more pigments, as they include the pigments used for photo protection. For every dataset, we linearly regressed the log transformed species richness and log transformed biovolume against pigment richness, and compared it with the theoretical results.

The empirical data we found differs in two aspects from our simulated data. First, our simulated data represent long-term effects of pigment richness on species richness and function under steady environmental conditions and it is not sure if the empirical data likewise represent such long-term effects. The field communities will have been exposed to environmental fluctuations and the experimental communities were observed during relatively short-term experiments (8-14 days). Second, the empirical data contain photosynthetic pigments as well as non-photosynthetic pigments, typically used for photoprotection. We therefore repeated our simulations, breaking them off after 10 days, hence simulating the time window typically adopted by experiments. We also included six non-photosynthetic active pigments, that absorb light, but do not contribute to growth (Bricaud *et al.*, 2004). Specifically we changed 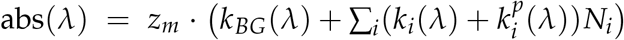 from equation 1, where 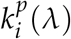 is the combined absorption spectra of the photo protective pigments. 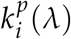 was generated similar to *k*_*i*_(*λ*).

## Results

### Effects of pigment richness on coexistence

We found that one can decompose the growth rate of a species as:

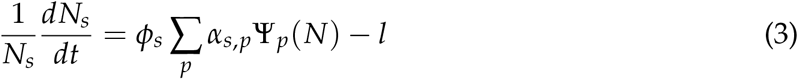

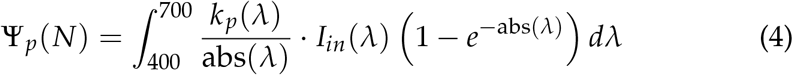

Where Ψ_*p*_(*N*) is the amount of light absorbed by pigment *p*. Stable and feasible coexistence, i.e. all species have non-zero equilibrium densities, would impose 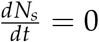 for all the species *s*. This yields a linear equation in the variables Ψ_*p*_(*N*). This equation can only have a solution if the number of species is at most equal to the number of pigments in the community. This finding proves the fact that species richness is limited by pigment richness for constant incoming light intensity (Meszéna *et al.*, 2006).

These new analytical results show that the number of pigments puts an upper bound on the number of coexisting species, but how many co-existing species can we on average expect at a given number of pigments? While equation 3 will always have a solution, this solution might not be ecologically possible. For example, all pigments should have a positive contribution to growth (Ψ_*p*_(*N*) > 0). In addition, pigments in nature often share peaks: Many pigments have maxima in the blue part of the spectrum (400*nm −* 450*nm*) such that the different Ψ_*p*_(*N*) are not independent functions (Fig. 1 and Stomp *et al.* (2007)).

Our simulations of species richness at various combinations of pigment richness and photosynthetic efficiencies show that the number of coexisting species is often lower than the number of pigments. Initial pigment richness increased average species richness from 1 to 2.2 (Fig. 2). When all pigments are present (initial pigment richness = 9), at most 4 species can coexist, but with low probability (≈ 2.5%). We repeated the analysis with several alternative assumptions on initial species richness (ranging from 1-25), the included pigments (including carotenoids with photo-protective purposes), the absorption spectrum of the pigments, the presence of pigments in species, the absorption spectrum of the background and the incoming light spectrum (see appendix 3). We also ran simulations in which absorption spectra of pigments where randomly generated and pigments were randomly distributed to species. All of these simulations lead to the same main conclusion: pigment richness is essential to allow coexistence but does mostly not allow the coexistence of more than two species in light limited phytoplankton communities.

**Figure 2:**
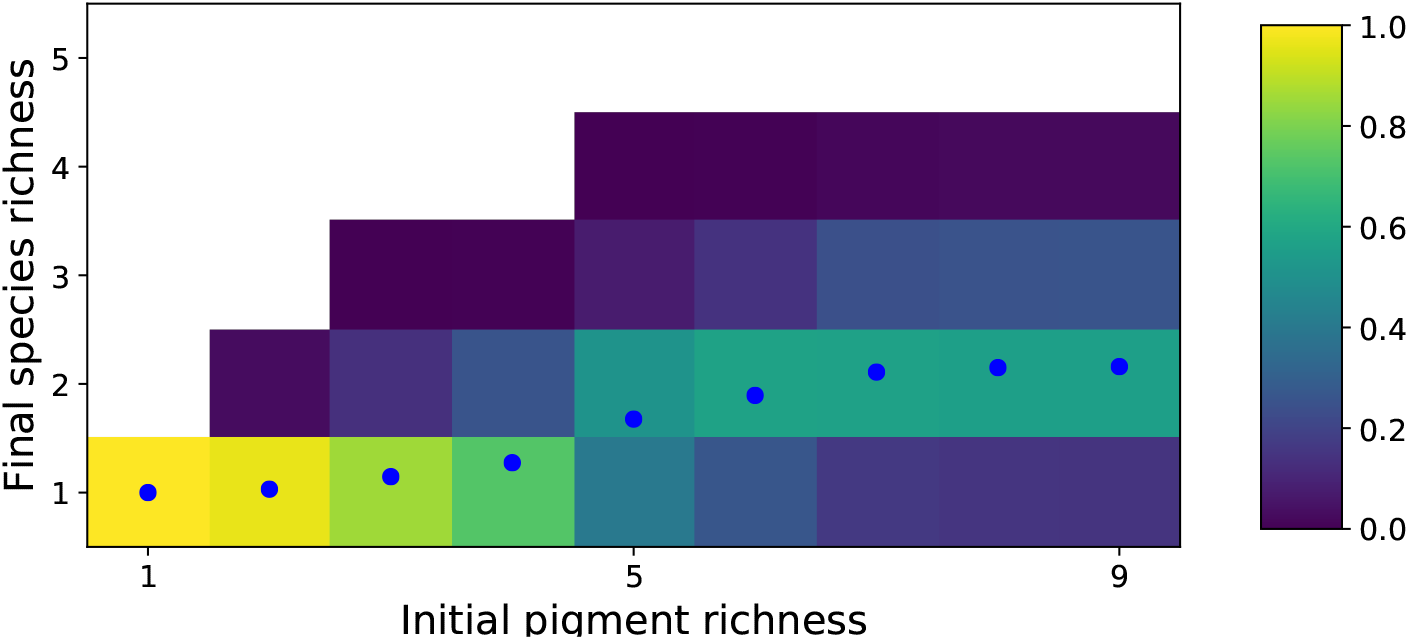
Final species richness depends only weakly on the initial number of pigments. Colours denote the probability associated with these species richness levels. Blue dots show the average of the final species richness.

We identified three reasons for this result, all related to how pigment richness affect niche 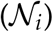 and fitness differences (*FD*_*i*_). First, pigment richness had a unimodal effect on 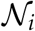 (Fig. 3 A). At low pigment richness, pigment richness increases 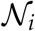, as it increases the probability species will be coloured differently, which stimulates light partitioning. As pigment richness increases, however, species tend to contain an increasing number of pigments, blurring differences in pigmentation, which reduces 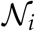. Second, pigment richness does not only affect 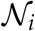, which benefits coexistence, but also increases differences among species’ 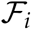, which hampers coexistence. Intermediate pigment richness lead to clear competitive dominant (high 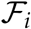) and subordinate (low 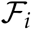) species (Fig 3 B). Communities with intermediate pigment richness will contain species with many pigments, that absorb most wavelengths and have high fitness, and species with few pigments, that absorb only part of the wavelength and have low fitness. The difference in these fitnesses leads to large 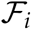, both positive and negative. As pigment richness increases all species become generalists and have high fitness, therefore 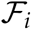 are closer to zero. Third, 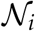 and 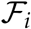 were located close to the origin of the 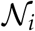 and 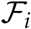 plane (Fig. 3 C), showing that different pigmentation types only offered limited opportunity for differentiation among species. This is explained by all pigmentation types containing chlorophyll a, which makes pigmentation more similar (Appendix S2, Van Den Hoek *et al.* (1995)), and different pigments having similar absorption spectra (Stomp *et al.*, 2007). Again, repetition of the simulations with altered parameters led to the same results, niche and fitness differences are located close to the origin in the 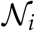 and 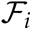 plane (Appendix 3).

**Figure 3:**
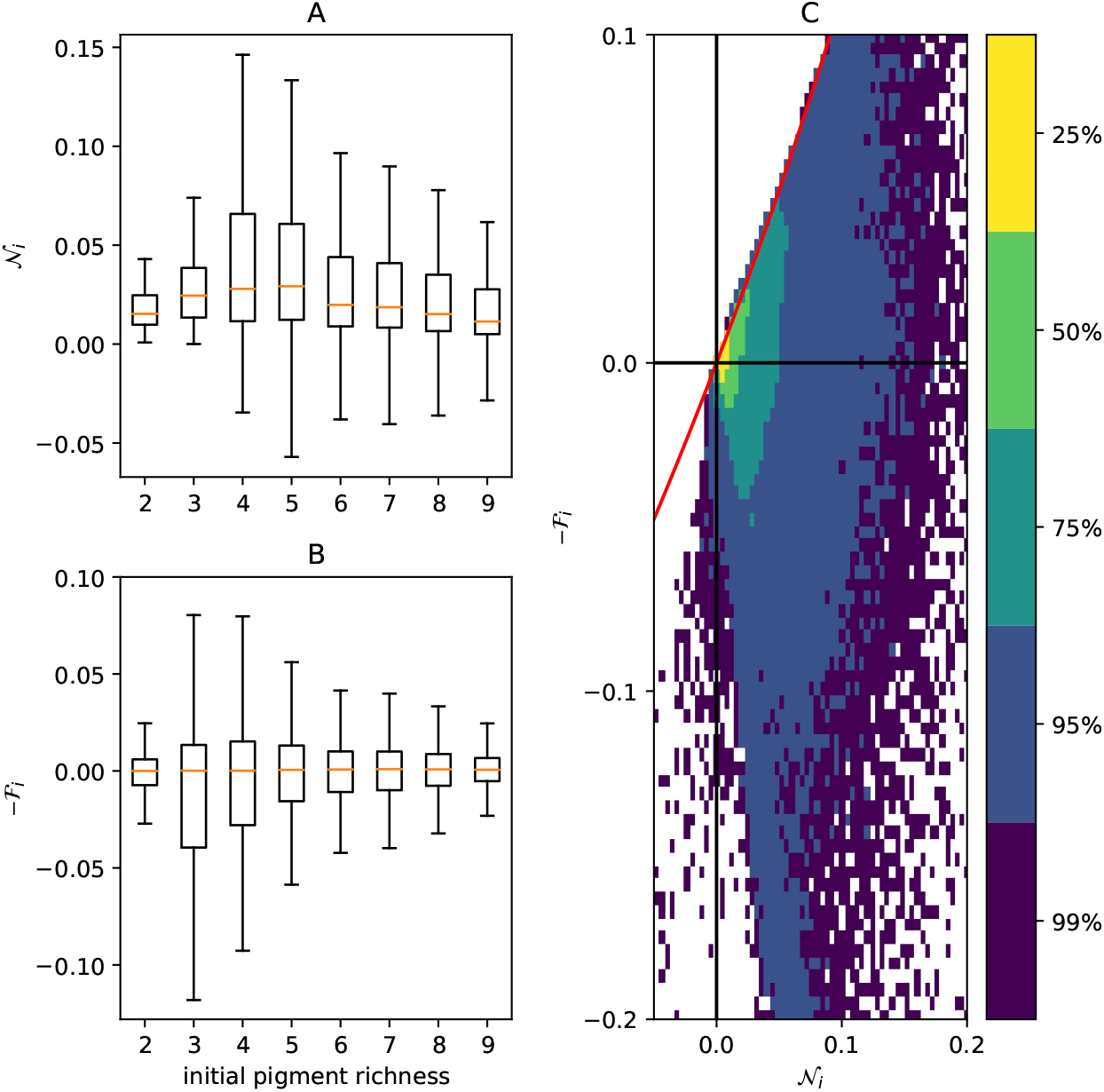
A: Pigment richness has a unimodal effect on 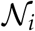. Initially multiple pigments allow niche differentiation by the use of different wave-length. However, this positive effect of pigment richness on species richness is limited to few pigments (up to about 4 pigments) and generates only small 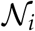 compared to empirical measurements in phytoplankton (Narwani *et al.*, 2013; Gallego *et al.*, 2019). At higher pigment richness the species tend to be more generalists and absorb multiple wavelengths, which reduces pigment richness. Shown are the 5, 25, 50, 75 and 95 percentiles of 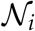 of all persisting species from all simulated communities. B: Similarly, pigment richness affects 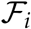. Communities with intermediate pigment richness are composed of species with many pigments, that have high fitness, and species with few pigments, that have low fitness, leading to both, strong negative and positive 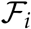. The species with strong negative 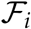 are most vulnerable to exclusion. Communities with low or high pigment richness are composed of species with few or many pigments, respectively. The species from these communities tend to have similar fitness, which leads to weak 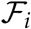. C: Two-dimensional histogram of 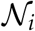 and 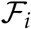 for all species in all communities, independent of their pigment richness. Most species have both 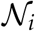 and 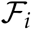 close to 0. Shown are the 25,50,75, 95 and 99 percentiles of the distribution, e.g. 25% of the species have 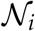 and 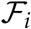 within the yellow area. Species below the red-line have a positive invasion growth rate, i.e. they are assumed to persist.

Contrary to our theoretical results, the compiled empirical data did suggest a strong positive effect of pigment richness on species richness (Fietz & Nicklisch, 2004; Estrada *et al.*, 2004; Striebel *et al.*, 2009). However, our model analyses and simulations represent the long-term outcome of species interactions in stable environmental conditions, while the empirical data describe experiments performed on much shorter time scales (approximately 10 days) and natural communities with unknown environmental conditions. Thus, the empirical data potentially describe communities that are in a transient state, heading towards lower levels of species richness. Our additional model simulations of short-term effects of pigment richness on species richness indeed suggest time scale to be an important explanation for the difference between our model results and the empirical data. These simulations showed that effects of initial pigment richness on species richness are indeed initially strong, but become weaker over time: pigment richness promoted short-term species richness (Fig. 4A).

**Figure 4:**
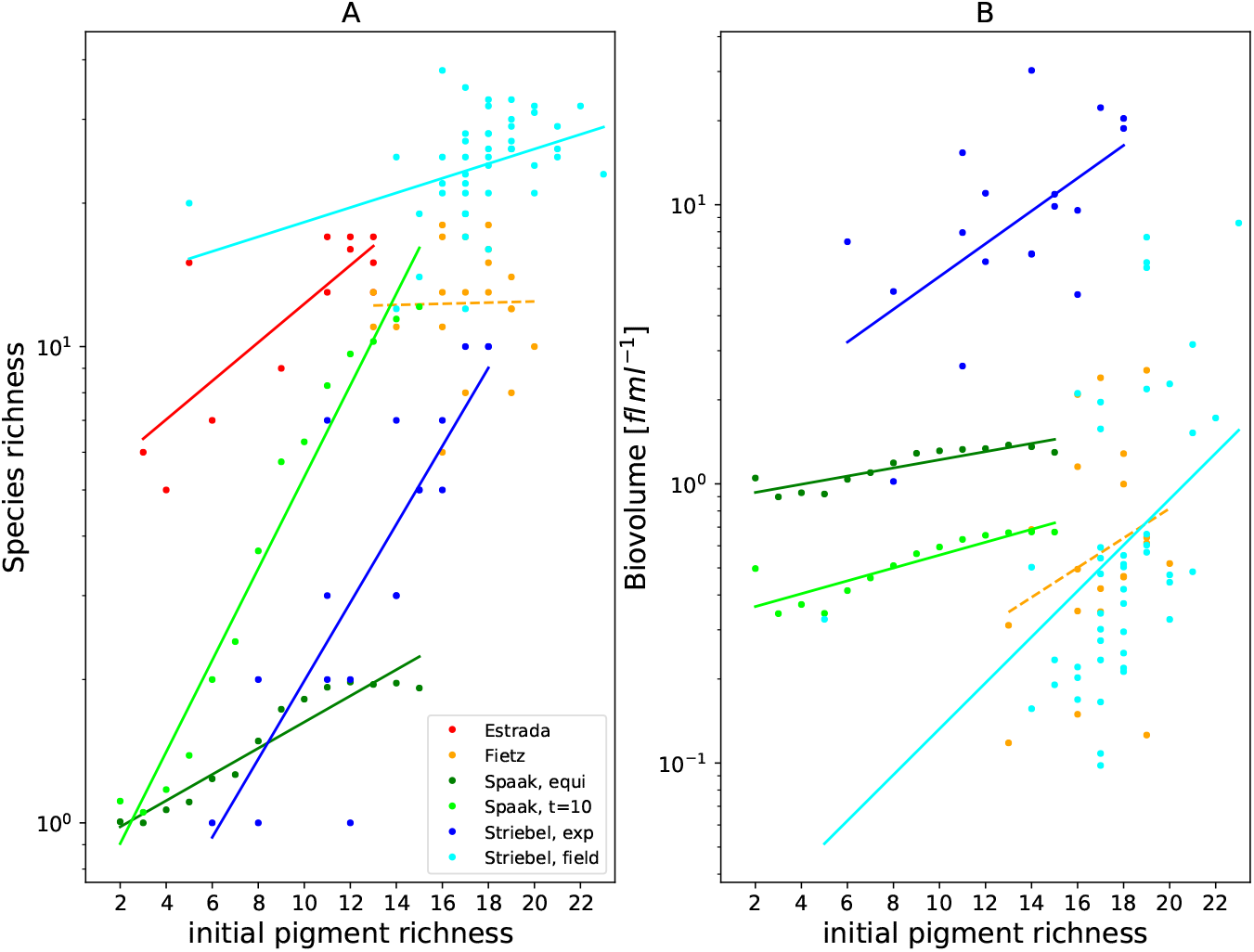
Empirical effect of initial pigment richness on species richness (A) and ecosystem function (biovolume B). A: Effects of pigment richness on species richness is initially strong (light green line), but weakens as the communities approach equilibrium densities (dark green line). Similarly, empirical data report a positive effect of pigment richness on species richness. This positive effect is stronger in experimental datasets (dark blue and red) than in natural communities (light blue and orange), presumably because natural communities are closer to equilibrium densities than short term experiments. B: The same observations hold true for the effects of pigment richness on ecosystem function, these are initially strong (slope of light green: 0.101 ± 0.012), but weaken over time (slope of dark green: 0.063 ± 0.007). Dashed lines indicate, that slope did not significantly differ from 0 (i.e. *p* > 0.05).

### Implications for ecosystem function

Our simulations show that ecosystem function in communities with high pigment richness is up to 60% higher than in communities with low pigment richness (total biomass, Fig. 4B). This increase of ecosystem function is mainly due to selection effect and not complementarity (Appendix 4). Complementarity is low, because light partitioning allows only limited niche differentiation. As found for species richness, the compiled empirical data did suggest a stronger positive effect (larger slope) of pigment richness on ecosystem function (Striebel *et al.*, 2009; Estrada *et al.*, 2004; Fietz & Nicklisch, 2004) (Fig. 4). Again, we additionally simulated short-term effects of pigment richness on ecosystem function (Fig. 4B, light green) and found a stronger effect of pigment richness on ecosystem function than for communities at equilibrium (dark green). The effects of initial pigment richness on function are initially strong but dampen with time.

## Discussion

In the past decade, ecology has evolved towards trait based approaches, as species interactions are governed through species traits and not through species identities (Litchman & Klausmeier, 2008; Mcgill *et al.*, 2006; Degen *et al.*, 2018). Trait based approaches have been used to explain species richness (Violle *et al.*, 2011; Best *et al.*, 2013), coexistence mechanisms such as niche and fitness differences (Kraft *et al.*, 2015; Narwani *et al.*, 2013; Gallego *et al.*, 2019), and ecosystem function (Gross *et al.*, 2017; Degen *et al.*, 2018). Here we investigated the effect of pigment richness on all these key properties of ecological communities using a mechanistic community model describing light limited phytoplankton. We found that pigment richness increases species richness from one to two species on average (Fig. 2), has a unimodal effect on niche and fitness differences, and increases ecosystem function by about 60%.

### Effects of pigment richness on species richness

Light has traditionally been considered as one resource until Stomp *et al.* (2004) showed that different wavelengths allow niche partitioning along the light spectrum, supporting coexistence. Our results confirm that this additional trait axis, representing pigmentation phenotypes, indeed allows introducing niche differences that allow coexistence of multiple species that would not have been possible if a single pigment would have been present (Stomp *et al.*, 2004, 2007; Passarge *et al.*, 2006; Huisman & Weissing, 1994). However, our results also show that this mechanism alone cannot explain robust coexistence of many more species than two.

One important reason is the fact that pigments not only stimulate niche differences, but they also create competitive dominant and sub-ordinate species, leading to large fitness differences, as they determine the total amount of absorbed photons 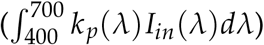. This result illustrates the effect of limiting similarity and the widespread occurrence of competitive exclusion despite trait differences (Meszéna *et al.*, 2006; Barabás *et al.*, 2012). While the light spectrum represents a continuous resource axis, along which an infinite number of species could specialise and persist, species that harvest too similar light colours will lead to the exclusion of the least fit. The fact that traits influence both niche and fitness differences has been pointed out for other traits in other systems as well (Mayfield & Levine, 2010). For example, Kraft *et al.* (2015) found that phenology of annual plants did not only promote niche differences but also inflated fitness differences. Gallego *et al.* (2019) found that a single trait, phytoplankton size, increases both niche and fitness differences. However, larger size differences between species does not promote coexistence.

Despite the extensive parameter range considered and the use of realistic pigmentation phenotypes, our analysis is a simplification of reality. In more complicated models, including additional factors, the effect of trait richness on species richness may be stronger. A first factor is individual-specific pigmentation. That is, the absorption spectrum of the same pigment may be different in different individuals from the same species, representing subtle but potentially important sources of inter- and intraspecific variation (Bricaud *et al.*, 2004; Six *et al.*, 2007; Bricaud *et al.*, 1995). Intraspecific variation of resource uptake traits can promote coexistence. For example, Hausch *et al.* (2018) recently showed that moderate intraspecific variation of bean weevils feeding on lentils and adzuki increased the resistance against invasion and invasion ability, thus promoting stable coexistence. However, theoretical results on annual plants obtained by Hart *et al.* (2016) suggest that initial intraspecific trait variation may also hamper coexistence. A second factor embodies trade-offs among traits. For example, species with less competitive pigmentation could have higher photosynthetic efficiencies, similar to larger phytoplankton species often having higher maximum growth rates (Langdon, 1988; Edwards *et al.*, 2015; Lavallée & Pick, 2002). Our assumption that all trait combinations were equally probable is therefore clearly a simplification of reality. A third factor is photoinhibition, which could facilitate coexistence, as it could turn our linear response of growth to light intensity into a non-linear response (Huisman & Weissing, 1994; Gerla *et al.*, 2011; Stomp *et al.*, 2007). This would introduce the potential for relative non-linearities, which can contribute to temporal niche differentiation in case of external light intensity fluctuations (Litchman & Klausmeier, 2001). Litchman and Klausmeier showed that relative non-linearities can sustain coexistence of light-limited phytoplankton communities (Litchman & Klausmeier, 2001). Another way in which light intensity fluctuations could affect coexistence is through storage effects (Chesson, 2003). However, only one of the three conditions required for the storage effects are met. Indeed, in our system, environmental conditions do covary with competition, however, the response of all species is similar to these co-variances and population growth is additive in environment and competition. By including a dormant stage, which is not sensitive to environmental fluctuations, Tredennick *et al.* (2017) showed that the storage effect leads to coexistence in a simple resource competition model. Including different phytoplankton life stages could serve as dormant stages in this model as well, and possibly promote a positive effect of pigment richness on species richness through storage effects.

### Implications for ecosystem function

Our model showed that pigment richness positively affected ecosystem function (Fig. 4). However, coexistence requirements often caused the exclusion of many species in communities with high initial trait richness. This led to levels of ecosystem function that are comparable to that of communities with lower initial trait richness in which all species persist. This mechanism reduces the positive effect of trait richness on ecosystem function. Therefore, our results show how the effects of trait richness on ecosystem function can be limited on time scales that are sufficiently long for community dynamics to emerge. The limited long-term positive effect of trait richness on ecosystem function we find is due to the selection effect. Our results show that higher initial trait richness increases the probability of having a highly productive species present in the community (Appendix S5). At the same time, complementarity is small in light limited phytoplankton communities (Appendix 4). Cadotte (2017) have found that complementarity was most prominent in species and trait-rich plant communities.

Positive effects of trait richness on species richness and ecosystem function have been found in many different study systems. For example, diversity of feeding traits in herbivorous marine amphipods increased coexistence (Best *et al.*, 2013), mouth size differences facilitated coexistence of two bacterivorous ciliates (Violle *et al.*, 2011), and phylogenetic diversity, species richness and average productivity were all found to correlate positively in savanna grasslands (Cadotte *et al.*, 2009).

However, many of these studies share two features that may lead to higher trait diversity effects on function than we report on here. First, these experiments often are too short to observe competitive exclusion, and so species can contribute to function before they go extinct. Our results also show that it can take many generations to observe competitive exclusion, over 200 generations in our system, which is longer than most-long term experiments. Similarly long times to competitive exclusion have been found for phytoplankton species competing for two limiting resources (Sakavara *et al.*, 2017). Second, available experimental studies often consider communities with relatively few species. Our results show that the trait richness effect on species richness and function is most pronounced when trait richness, and therefore species richness, is relatively low (between 2 and 7 pigments).

Effects of trait diversity on ecosystem function of grasslands have been found to intensify with time (Reich *et al.*, 2012). This temporal intensification has been explained by reciprocal feedbacks between community composition and environmental conditions such as on soil nitrogen availability (Reich *et al.*, 2012). Our results show that, in absence of such feedbacks and when given enough time, effects of biodiversity on ecosystem function can actually weaken with time. That is, packing a more diverse set of species into our model system, light-limited phytoplankton, caused benefits for ecosystem function that were short-lived. These findings highlight the need to account for coexistence requirements when estimating the long-term benefits of biodiversity for ecosystem function (De Laender *et al.*, 2016; Bannar-Martin *et al.*, 2018).

## Acknowledgements

We thank J. Huisman for comments on earlier versions of this manuscript and on the collection of pigmentation data. We thank M.Striebel and S.Fietz for providing data. This research used resources of the Plateforme Technologique de Calcul Intensif (PTCI) located at the University of Namur (http://www.ptci.unamur.be), Belgium, which is supported by the FRS-FNRS under convention No 2.4520.11. The PTCI is member of the Consortium des Équipements de Calcul Intensif (CÉCI) (http://www.ceci-hpc.be). F.D.L. received support from grants of the University of Namur (FSR Impulsionnel 48454E1), and the Fund for Scientific Research, FNRS (PDR T.0048.16).

## Data accessibility statement

In case of acceptance we will put all code to perform this study on our git repository. No new experimental or field data was used in this study.

## Notes

### Competing Interest Statement

The authors have declared no competing interest.

