## Appendix for "Unimodal effects of pigment richness on niche and fitness differences explain species richness and ecosystem function in light-limited phytoplankton communities"

### 1 Spectrum model

The equation given by Stomp *et al.* (2004) is equivalent to:

$$\frac{1}{N_s}\frac{dN_s}{dt} = \frac{\phi_s}{z_m}\int_0^{z_m}\int_{400}^{700} k_s(\lambda) I_\lambda(z) d\lambda\, dz - l_s \tag{1}$$

$$= \frac{\phi_s}{z_m}\int_{400}^{700}\int_0^{z_m} k_s(\lambda) I_\lambda(z) dz\, d\lambda - l_s \tag{2}$$

$$= \frac{\phi_s}{z_m}\int_{400}^{700} k_s(\lambda)\int_0^{z_m} I_\lambda(z) dz\, d\lambda - l_s \tag{3}$$

$$= \frac{\phi_s}{z_m}\int_{400}^{700} k_s(\lambda)\int_0^{z_m} I_{in}(\lambda) e^{-z\left(\sum_i \frac{N_i}{z_m} k_i(\lambda) + k_{BG}\right)} dz\, d\lambda - l_s \tag{4}$$

$$= \frac{\phi_s}{z_m}\int_{400}^{700} k_s(\lambda) I_{in}(\lambda)\left[\frac{e^{-z\left(\sum_i \frac{N_i}{z_m} k_i(\lambda) + k_{BG}\right)}}{-\left(\sum_i \frac{N_i}{z_m} k_i(\lambda) + k_{BG}\right)}\right]_0^{z_m} d\lambda - l_s \tag{5}$$

$$= \phi_s \int_{400}^{700} \frac{k_s(\lambda) I_{in}(\lambda)\left(1 - e^{-(\sum_i N_i k_i(\lambda) + z_m k_{BG})}\right)}{\sum_i N_i k_i(\lambda) + z_m k_{BG}} d\lambda - l_s \tag{6}$$

$$= \phi_s \int_{400}^{700} \frac{k_s(\lambda)}{\mathrm{abs}(\lambda)} \cdot I_{in}(\lambda)\left(1 - e^{-\mathrm{abs}(\lambda)}\right) d\lambda - l_s \tag{7}$$

Where we introduced the species independent variable $\mathrm{abs}(\lambda) = \sum_i N_i k_i(\lambda) + z_m k_{BG}$ for convenience.

#### 2 Pigment table

| Pigment | Chl a | Chl b | Chl c | Beta Carotene | Peridinin | Fucoxanthin | PCB | PEB | PUB |
|---|---|---|---|---|---|---|---|---|---|
| Heterokontaphyta 1 | 1 | | | | | | | | |
| Heterokontaphyta 2 | 1 | | 0.2 | 0.3 | | | | | |
| Heterokontaphyta 3 | 1 | | 0.2 | 0.3 | | 0.5 | | | |
| Haptophyta | 1 | | 0.2 | 0.3 | | 0.5 | | | |
| Prochlorophyta | 1 | 0.2 | | 0.3 | | | | | |
| Euglenophyta | 1 | 0.2 | | 0.3 | | | | | |
| Chlorarachniophyta | 1 | 0.2 | | | | | | | |
| Chlorophyta | 1 | 0.2 | | 0.3 | | | | | |
| Dinophyta I | 1 | | 0.2 | 0.3 | 0.5 | | | | |
| Glaucophyta | 1 | | | 0.3 | | | 1 | | |
| Rhodophyta | 1 | | | 0.3 | | | 0.25 | 1 | |
| Cryptophyta | 1 | | 0.2 | | | | 0.25 | 1 | |
| Cyanophyta 1 | 1 | | | | | | 1 | | |
| Cyanophyta 2 | 1 | | | | | | 0.25 | 1 | |
| Cyanophyta 3 | 1 | | | | | | 0.25 | 0.25 | 1 |

Table 1: Abberations for pigments: Chl: Chlorophyll, PCB: Phycocyanobilin, PEB: Pyhcoerithrobilin, PUB: Phycourobilin. Shown are only the main pigments according to Van Den Hoek *et al.* (1995) and Six *et al.* (2007). The concentrations of the pigments are taken from Bricaud *et al.* (2004).

#### 3 Additional simulations

For the simulation chosen in the main text we took quite strong assumptions, which will potentially not hold in nature. We made a large number of different simulations, none of which changed the general result obtained that trait richness hardly increases species richness. Additionally in all cases the positive effects of trait richness on ecosystem function were more pronounced for short term experiments, in which hardly any species went extinct, than for long term experiments.

These simulations included slight variation of pigment absorption spectra per species, back ground absorption and different fluctuations of incoming light. As an additional example we show the case where the pigment absorption spectra varied up to 5% between species and the background absorption resembles that of a coastal region Stomp *et al.* (2007) with and without fluctuation of incoming light.

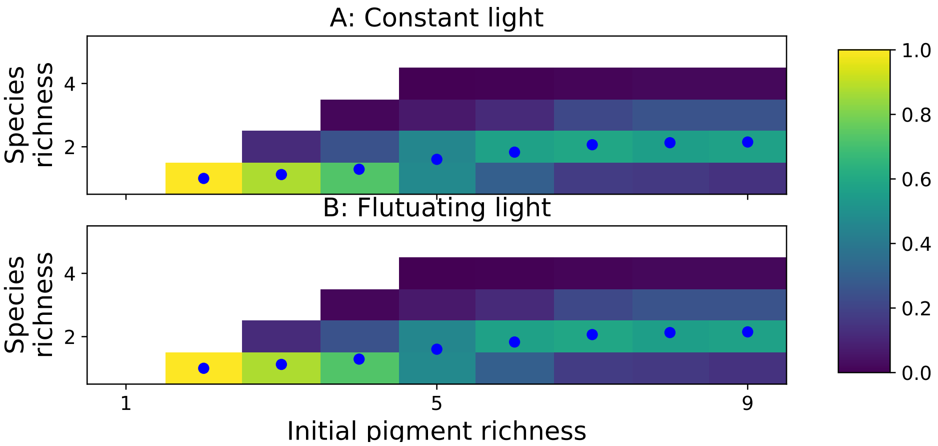


Figure 1: Final species richness depends only weakly on the initial number of pigments at constant (A) and fluctuating (B) incoming light. Colours denote the probability associated with these species richness levels. Blue dots show the average of the final speices richness. In this simulation we included a background absorption of a coastal region Stomp *et al.* (2007). Furthermore the interspecific variation of the absorption spectra of the pigment was assumed to be 5%.

### 4 Selection and complementarity

Most species have few different pigments ($\leq$ 3) (See table 1), hence a community with high initial pigment richness is likely to also have high initial species richness. High initial species richness, however, implies higher probability of having a highly productive species present in the ecosystem. We compute selection, complementarity and relative yield total as described in Hector *et al.* (2001).

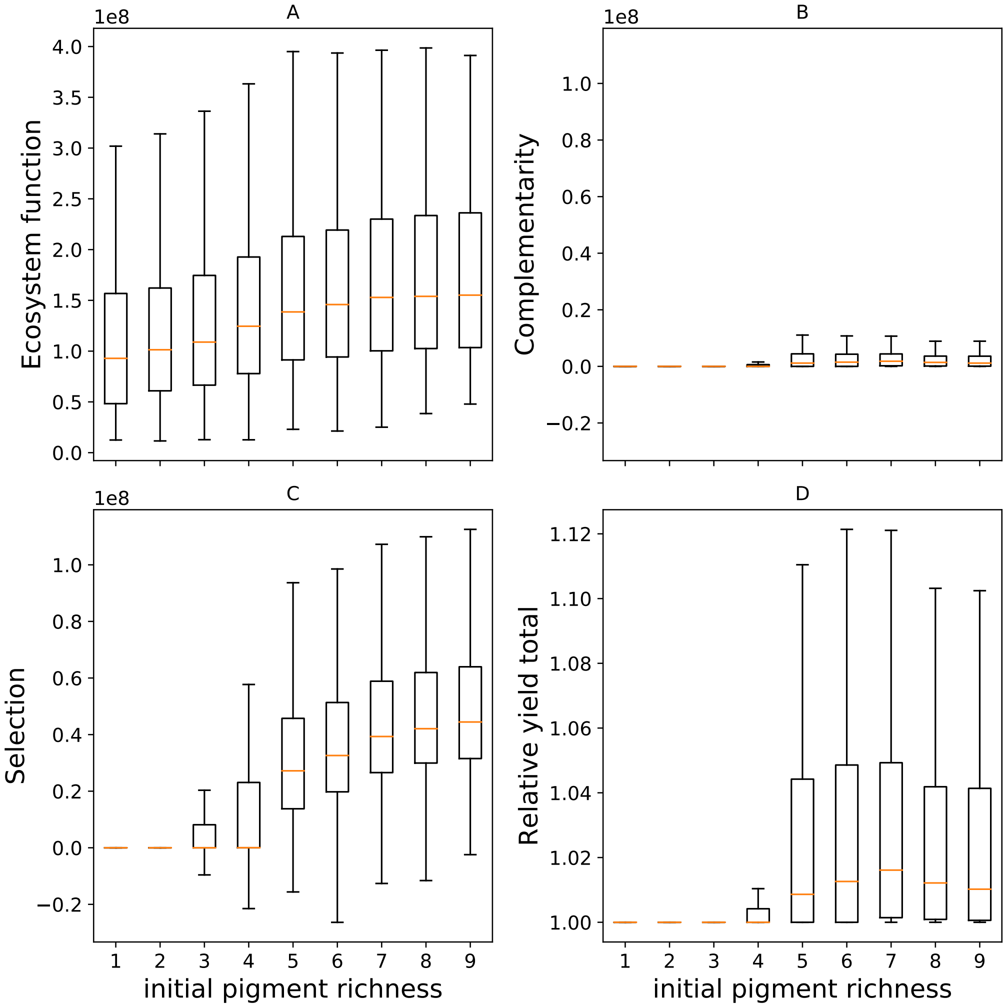


Figure 2: The positive effect of pigment richness on ecosystem function (A) can be decomposed into complementarity, which is associated with resource partitioning, (B) and selection effect, which is associated with dominance of productive species (C). Initial pigment richness increases both, complementarity and selection effect, however, the selection effect is an order or magnitude stronger than the complementarity effect. D: Initial pigment richness increases relative total yield by 1-2%, which should be considered as minor. That is communities with high pigment richness do not yield much more, than the most productive monoculture species. Shown are 5%, 25%, 50%, 75% and 95% percentiles.
